# Is the Stalk of the SARS-CoV-2 Spike Protein Druggable?

**DOI:** 10.1101/2022.10.06.511069

**Authors:** Ludovico Pipitò, Christopher A. Reynolds, Giuseppe Deganutti

**Affiliations:** Centre for Sport, Exercise, and Life Sciences, Coventry University, CV1 5FB, UK

**Author notes:** To whom correspondence should be addressed, GD.

## Abstract

The SARS-CoV-2 virus spike protein (SP) is the vector of the virus infectivity. The high propensity to mutate in key regions responsible for the recognition of the human angiotensin-converting enzyme 2 (hACE2) or the antibodies produced by the immune system following infection or vaccination makes subunit 1 of the SP a difficult to target and, to date, efforts have not delivered any ACE2 binding inhibitor yet. The inherent flexibility of the stalk region, within subunit S2, is key to SARS-CoV-2 high infectivity because it facilitates the receptor binding domain encounter with ACE2. Thus, it could be a valuable therapeutic target. By employing a fragment-based strategy, we computationally studied the druggability of the conserved part of the SP stalk by means of an integrated approach that combines molecular docking with high-throughput molecular dynamics simulations. Our results suggest that the druggability of the stalk is challenging and provide the structural basis for such difficulty.

## Introduction

The severe acute respiratory syndrome coronavirus type 2 (SARS-CoV-2) pandemic that was first identified in 2019 in the city of Wuhan ^1^ continues to raise concerns amongst governments and the scientific community with almost 530 million cases around the world with more than 6.2 million certified deaths (WHO dashboard, 08 June 2022, https://covid19.who.int). SARS-CoV-2 shows a strong affinity for the human angiotensinconverting-enzyme 2 (ACE2) receptor, a type 1 transmembrane protein responsible for the extracellular conversion of the angiotensin hormone into angiotensin II ^2^ through the SARS-CoV-2 spike protein (SP). The SP is a highly glycosylated trimeric structure, common amongst the coronaviridae family ^3^, which is constituted by two main units named S1 and S2. While S1 is responsible for molecular recognition of ACE2, S2 is paramount to structural stability and orientation of the whole SP and membrane fusion to deliver the viral genome ^4^. The efforts of the scientific community were dedicated to promptly developing vaccines or a variety of small molecules^5–8^, specifically designed to bind and neutralize the area on S1 responsible for ACE2 binding, namely the receptor-binding domain (RBD).

In 2020, the term variants of concern (VOC) was introduced to describe new SARS-CoV-2 strains which differentiated from the original SARS-CoV-2 wild type (WT) through a series of mutations, mainly on the RBD, which cause drastic changes in transmissibility and pathogenicity ^9–11^. The B.1.617.2 strain (Delta variant) was identified in India by January 2021 and spread rapidly across the globe^12^, overcoming the WT in a short amount of time. n November 2021 the B.1.1.529 (Omicron variant) became the dominant VOC over the Delta^13^. Among the SARS-CoV-2 VOCs, major preoccupations regarded those strains that carried important mutations and deletions, especially on the RBD ^14^. VOC has important RBD mutations: B.1.1.7 (Alpha), carries E484K, N501Y, D614G, P681H; B.1.351 (Beta) carries K417N, E484K, N501Y, D614G, A701V; P1 (Gamma) carries K417T, E484K, N501Y, D614G, H655Y; B.1.617.2 (Delta) carries L452R, T478K, D614G, P681R ^15^. Concerns among the scientific community have risen due to their potential to elude the immune system and overcome vaccine protection ^16,17^ despite showing an overall similarity between variants, which diverged only in terms of flexibility of SP. More recently, a new B.1.1.529 (Omicron) VOC ^18^ carrying N440K, G446S, S477N, 118 T478K, E484A, Q493R, G496S, Q498R, N501Y, and Y505H mutations, and its lineages became predominant over the Delta variant, possibly due to a more rapid entry or different mechanism ^19–21^, an enhanced ability to evade the immune system^22–24^, and its increased affinity for ACE2 ^25–27^ although showing a milder pathogenic impact ^28^. New VOCs are expected to pose a new threat should they become widespread ^22,29^ and further studies should follow to evaluate the potential risk of new mutations.

The vaccine technology developed so far is designed to specifically target RBD, where the majority of the mutation occurred, increasing the risk for antibody inefficacy ^30–32^. The potential loss of efficacy against the Omicron variant was attenuated by a loss of replication and lethality power, probably due to Omicron’s inefficiency to exploit the cellular transmembrane protease serine 2 (TMPRSS2) ^28^. However, with the continuous viral diffusion, the likelihood of new mutations remains critical and new variant-specific vaccines need to be developed regularly to keep up with the rate of mutation ^33^. While S1 and RBD are the SP domains most prone to mutation, S2 has a higher level of conservation among the coronaviridae family ^34^. The only S2 mutations identified so far are N764K, D796Y, N856K, Q954H, N969K, and L981F. Residues I921, S980, V1187, F1218, and I1219 **(Figure 1)**, conserved in both the Delta and Omicron strains, are pivotal residues that confer increased flexibility to the stalk^35,36^. Interestingly, the region between residues L1145-L1186 (Conserved Region 1, **Figure1**), and between E1188-W1217 (Conserved Region 2, **Figure1**) contain Loop 1 and Loop2 that contribute to the S1 domain flexibility^37^, are conserved amongst all the VOC, and exhibit specific highly-conserved sequences^38^. Molecular dynamics (MD) simulations highlighted unexpected flexibility of the SP stalk ^36^, which has been proposed as paramount for ACE2 binding and infectivity ^39^. In principle, a small molecule able to target the stalk region would be effective on all the VOC by impairing the flexibility of the SP, thus the infectivity of SARS-CoV-2. For this reason, we integrated molecular docking and molecular dynamics (cMD) simulations to study the druggability of the conserved S2 stalk region E1144-R1185 **(Figure 1B)** and its potential as a drug target. Possible cryptic binding sites were sampled using a mixed MD (mixMD) approach^31,40,41^ to evaluate the accessibility of the stalk in the presence of the branched glycans on the SP surface. We docked a small library of optimized fragments^32^ to the SP stalk and performed hundreds of high-throughput post-docking cMD simulations combined with binding free-energy calculations to determine if small molecules can target this important SARS-CoV-2 protein domain.

**Figure 1.**
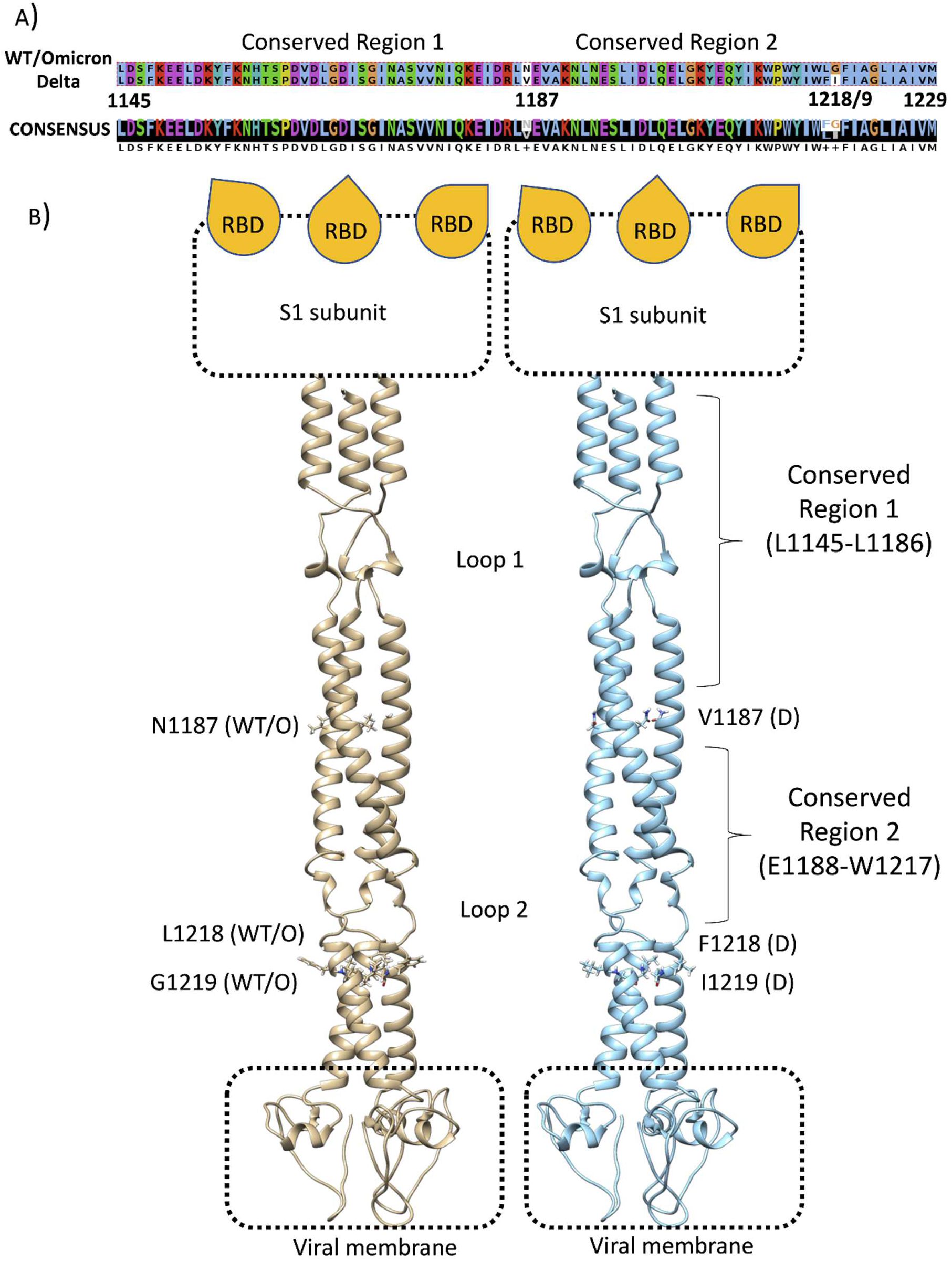
T. A) Sequence alignment between the conserved SP stalk region of the wild type (WT) or Omicron and Delta variants. **B)** Structural comparison between WT or Omicron (tan ribbon) and Delta S2 region (cyan). The area between residues L1145-W1217 is almost identical between the strains and could represent a therapeutic target. Glycans were removed for clarity; the S1 subunit with the three receptor binding domains (RBDs)and the viral, membrane are schematically represented.

## Methods

### General Workflow

MD simulations and molecular docking were combined in a computational pipeline (**Figure 2)** aimed to discern potential fragments able to overcome the steric barrier provided by glycosylation and engage the SP stalk in specific interactions. Initial MD simulations of the stalk prepared the structure for molecular docking, while mixMD sampled the accessibility of potential pockets. Molecular docking followed by high throughput post docking MD simulations discerned the stability of the predicted poses, narrowing the number of putative binders to five, which were further evaluated in further, extended MD and mixMD simulations.

**Figure 2.**
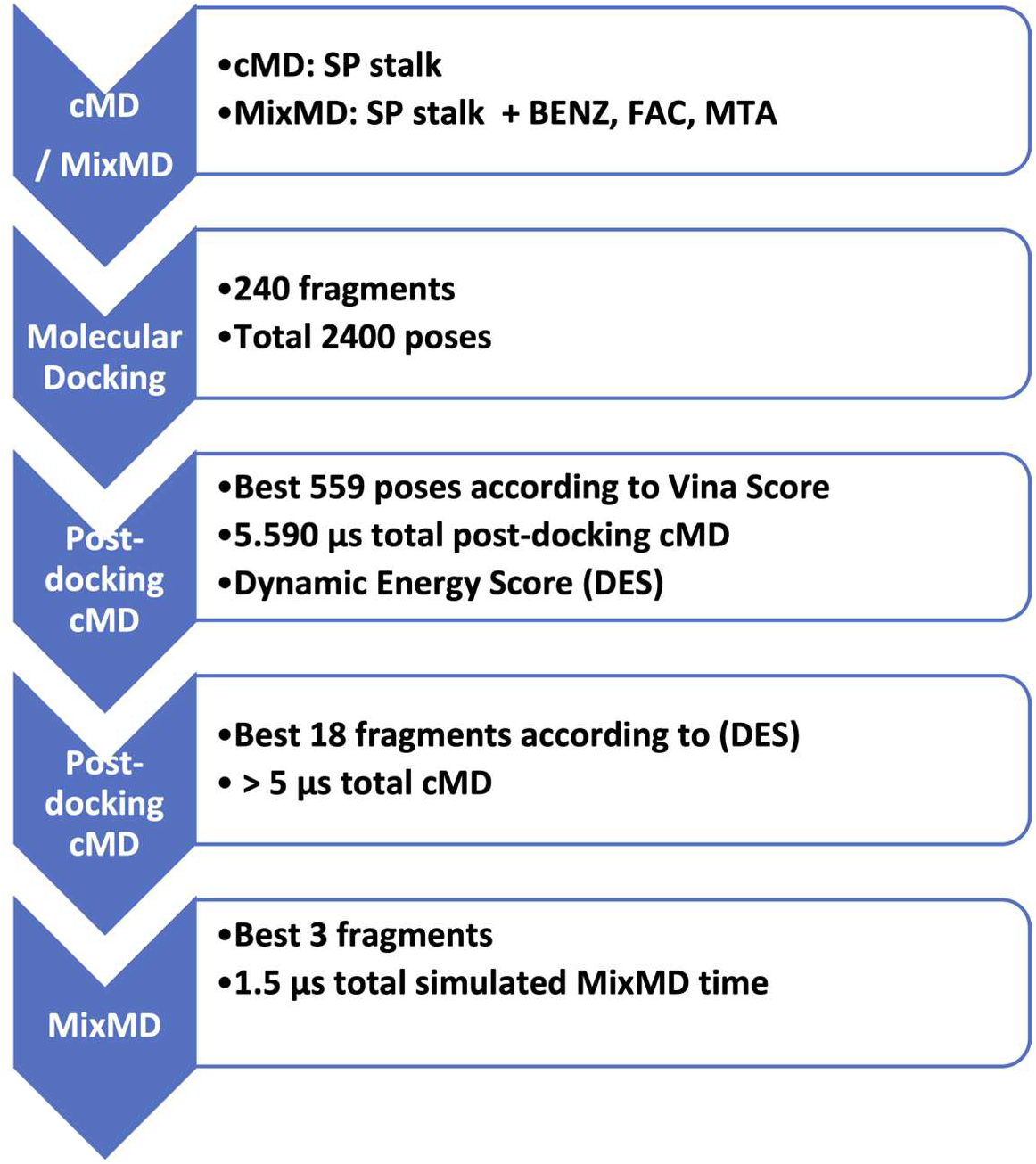
Computational workflow of the study. Preliminary classic molecular dynamics (cMD) and mixed MD (mixMD) were performed on the spike protein stalk (SP); an MD-extracted conformation of the SP stalk was used to dock 240 molecular fragments. The best 560 poses according to Vina score were subjected to 10 ns of cMD ed evaluated according to the dynamic energy score (DES, Equation 1). The best 5 fragments according to DES were finally further simulated through cMD and mixMD.

### Classic MD of the spike protein stalk

The SP stalk was prepared using CHARMM36 ^42,43^. The fully glycosylated SP model was retrieved from the CHARMM-GUI repository (https://charmm-gui.org/?doc=archive&lib=covid19), ^44^, and subsequentially trimmed from residue E1144-W1214, keeping the glycans in their original position at their original length. Hydrogen atoms in the S2 domain were added by Propka ^45^ at a simulated pH of 7.0, while structural integrity was checked through HTMD^46^, visually inspected, and patched manually through VMD ^47^ according to previous structural knowledge ^4^. Each system was solvated with TIP3P water molecules^48^ added to the simulation box considering a 15 Å padding in every direction by Solvate plugin 1.5 (http://www.ks.uiuc.edu/Research/vmd/plugins/solvate/). The charge neutrality was achieved by adding Na^+^/Cl^-^to the concentration of 0.150 M using Autoionize plugin 1.3 (http://www.ks.uiuc.edu/Research/vmd/plugins/autoionize/). The initial geometry and internal energy were optimized using the conjugate gradient algorithm by ACEMD^49^ to eliminate possible clashes and optimize atomic distances. The equilibration was achieved in isothermal-isobaric conditions (NPT) using the Berendsen barostat^50^ (target pressure 1 atm) and the Langevin thermostat^51^ (target temperature 300 K) with low damping of 1 ps^-1^. During the 4 ns equilibration, a positional restraint of 1 kcal/ mol Å^2^ was applied on the alpha carbons for the first 3 ns, and on protein side chains for the first 2 ns.

### Fragments preparation and molecular docking

A set of 240 optimized molecular fragments (**Table S1**) from X-ray complexes^32^, the SpotXplorer database, was converted to 3D conformers through the RDkit module^52^, protonated at pH 7.4 with Chimera^53^, and energy minimized with RDkit. Each fragment was docked using Autodock Vina^54,55^, to the Conserved Region 1 of the SP stalk using the structure from the last frame of the cMD equilibration (**Figure S1)** and the residue V1164 as the center of a grid with a 46 Å side length, for a broad exploration of the stalk surface. For each fragment, ten poses were ranked according to the docking score. Poses in contact with glycans or outside the conserved region were discarded. The rationale behind our selection was to narrow our list of fragments to those able to specifically target protein residues of the stalk. Poses away from the density maps produced by benzene (BENZ), formic acid (FAC), and methylamine (MTA) (see below) or not engaging simultaneously with at least two monomers, were also excluded.

### Post-docking classic molecular dynamics (cMD)

The best 559 docking complexes were subjected to 10 ns of post docking cMD. Initial CGenFF force field^56,57^ topology and parameter files of molecular fragments were obtained from the CGenFF software. Restrained electrostatic potential (RESP) charges were calculated with AmberTools20^58^ after geometry optimization through Gaussian09 at the HF/6-31G* level of theory^59^. Each complex was prepared for cMD, equilibrated, and simulated as reported below. For each simulation, similarly to Sabbadin *et al*^60^, we calculated the dynamic energy score (DES, **Equation 1**), which is the sum, over all the MD frames, of the ratio between the generalized born surface area GBSA binding energy and fragment root mean square deviation (RMSD) to the initial docking pose, using AmberTools20 and VMD.

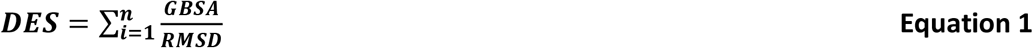

We excluded all the complexes with an average RMSD < 10 Å, and RMSD standard deviation < 5 Å, and ranked them according to the best final score. We then chose the five best fragments, which were visually inspected to avoid important interactions with glycans. These candidates were then further evaluated with 500 ns of cMD or mixMD (see below).

### Mixed Molecular Dynamics (MixMD)

Fragments 1-3 and three common molecular probes such as benzene (BENZ), formic acid (FAC), and methylamine (MTA) were used for mixMD^31,40^ to explore the accessibility of the stalk region to both very low (BENZ, FAC, MTA) and intermediate (fragments 1-3) molecular weight molecules, as well as possible cryptic binding sites. MixMD systems were prepared using PACKMOL ^61^, setting a minimum distance of 4 Å between each component to avoid clashes and secure a broad placement of the cosolvent molecules. An adequate number of cosolvent molecules were introduced to reach a virtual concentration of 5% w/w. The systems were then solvated, neutralized, equilibrated, and simulated as reported above. Density maps were computed using the Volmap VMD plugin (https://www.ks.uiuc.edu/Research/vmd/plugins/volmapgui/) setting a grid of 0.5 Å while Solvent accessible surface area (SASA) were estimated using vmdICE ^62^ and Chimera.

## Results

### The flexible loops promote stalk flexibility

Our investigation focused on the SP stalk Conserved Region 1, between residues L1145 and L1186, which is conserved and less glycosylated than other SP domains. During preliminary cMD simulations of the stalk (**Figure 3)**, the trimer maintained a stable quaternary structure in the region L1154-L1166, while displaying the highest flexibility at the level of the N-terminus (residues 1144-1156) and C-terminus (residues 1202-1214). The flexible Loop 1 (residues 1160-1170) was characterized by intermediate flexibility. The high RMSF (**Figure 3A)** of the N- and C- *termini* is ascribable to the artificial cut necessary to isolate the stalk region from the rest of the SP, which created unnatural protein ends. For this reason, we excluded the terminal four N-terminal amino acids for the successive docking studies. Computational studies have suggested high flexibility of the SP stalk due to the presence of two unstructured knees ^63^. Our simulations displayed similar behavior, where the bending of the stalk was aided by the opening of transitory pockets within the flexible loops, which temporarily moved away more than 20 Å from each other (**Figure 3B**) with an angle of about 133° between the alfa carbons of P1143-V1164-I1172 (**Figure S1)**.

**Figure 3.**
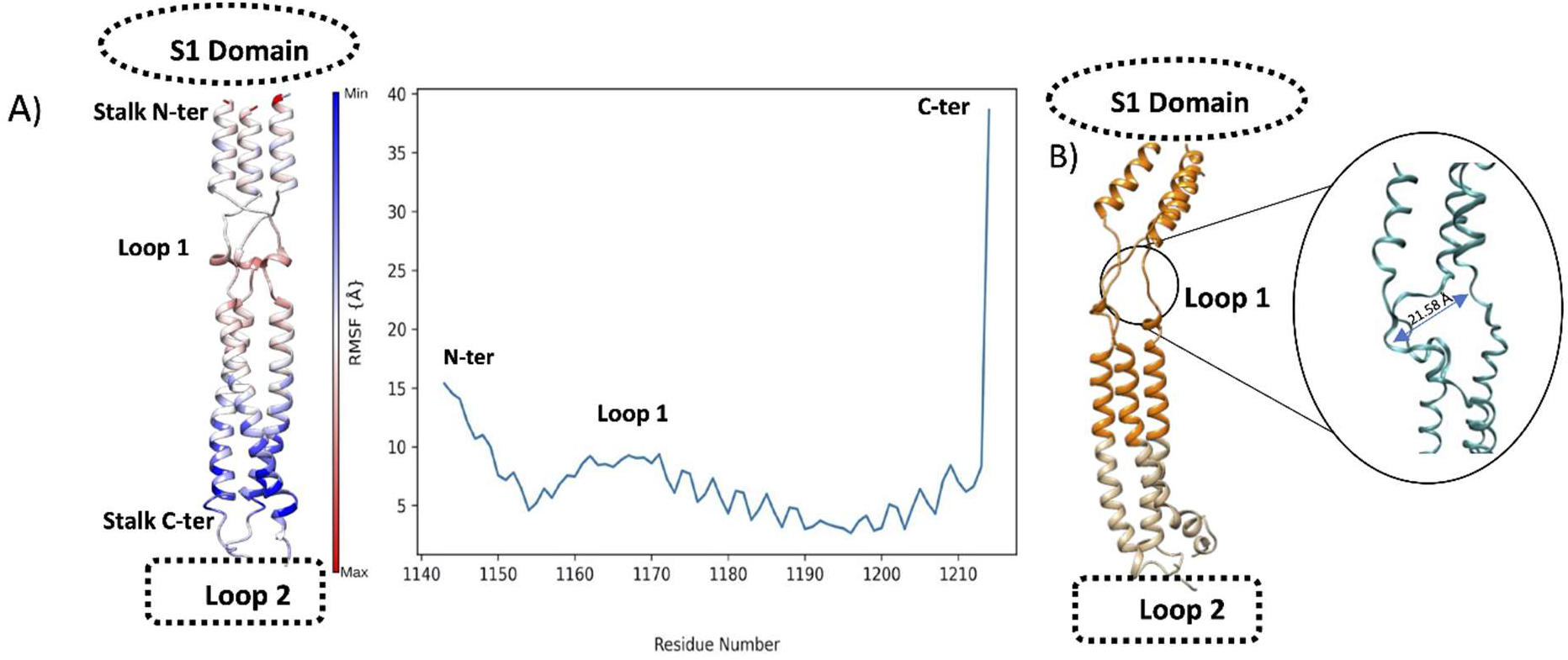
**A)** Root mean square fluctuation (RMSF) of the SP stalk during cMD simulations. The RMSF of each residue is mapped on the structure (left panel) and color coded according to the value (flexible residues are red) plotted; RMSF are also plotted on the sequence (right panel); **B)** the SP stalk Conserved region 1 (orange) is subjected to high flexibility during cMD simulations at the level of Loop 1; the position of the S1 domain and the Loop 2 is reported for reference.

### The conserved region of the stalk is accessible to solvent and small fragments

Preliminary MixMD simulations were employed to assess the accessibility of the Conserved Region 1 **(Figure S2)**. The probes benzene (BENZ), formic acid (FAC), and methylamine (MTA) indicated accessible sections across the stalk. BENZ sampled possible hydrophobic pockets on both the N- (close to K1149) and C-terminal (close to E1195) of the stalk. The latter was explored also by FAC, alongside a further interhelical volume in the proximity of E1182. MTA, and to a lesser extent BENZ, weakly interacted with the stalk at the level of the flexible loops’ residues V1164-S1170. These results indicate that the glycosylation of the stalk efficiently protects the SP, although some small areas are accessible for potential binding. Interestingly, molecular probes were able to intercalate within the three stalk helices, indicating possible pockets. SASA analysis indicated more solvent-exposed sites at the N-terminal of the stalk, below the connection with S1, and below the flexible Loop 1 (**Figure S3)** in accordance with mixMD results.

### Molecular docking and high throughput post-docking cMD predict interactions with Loop 1

Molecular docking of the optimized small library of fragments, performed on the fully glycosylated stalk, produced binding poses with scores ranging from −6.7 kcal/mol to −2.3 kcal/mol (**Table S2**) with limited convergences with density maps obtained from MixMD simulations (**Figure S2**). We discarded the last half of the poses (poses 1201-2400) as their docking score was lower than the arbitrary value of −4.5 kcal/mol and all the poses that did not interact with at least two monomers of the stalk or that made contacts with the glycans. The remaining 559 poses were then simulated during 10 ns of post-docking MD simulations, for a total MD sampling of almost 5.6 μs, in a fully hydrated and flexible environment. We computed the DES (**Equation 1**), which considers both the generalized born surface area (GBSA) binding energy and the root mean square deviation (RMSD) to the initial docking pose, for each fragment (**Table S3**). After visually inspecting the resulting best poses according to the DES, we kept only those closest to the Conserved Region 1 and not in contact with any glycan residue, retaining 18 fragments (**Table S4**), that were subjected to a further 500 ns cMD simulation.

### Longer post-docking MD simulations rebut molecular docking predictions

Disappointedly, 14 out of the 18 fragments were completely displaced in the first 100 ns of the extended simulations (**Table S4**). Rotations, openings, and closures of the flexible Loop 1 rapidly disentangled the fragments, causing the rapid displacement of most of the fragments. The remaining five compounds (fragments **1-5**, **Figure 4**, **Table 1, Video S1**) initially interacted with Loop 1, in correspondence with residues N1158-S1170, before moving away from the initial position. Overall, all the fragments changed orientations towards and formed alternative interactions before being displaced after less than 300 ns (**Figure 4E-I**). Planar compounds, predicted by docking in correspondence with the volumes identified by the initial mixMD as possible pockets, displayed better interactions with Loop 1 residues N1158-S1170. **1-3** resided at the center of the trimer for more than 100 ns, before moving upward towards the N-terminal through the three chains and dissociating; **4** and **5** resided mainly on the C-terminal end of Loop 1 before unbinding through a temporary tunnel formed between the stalk chains.

**Figure 4.**
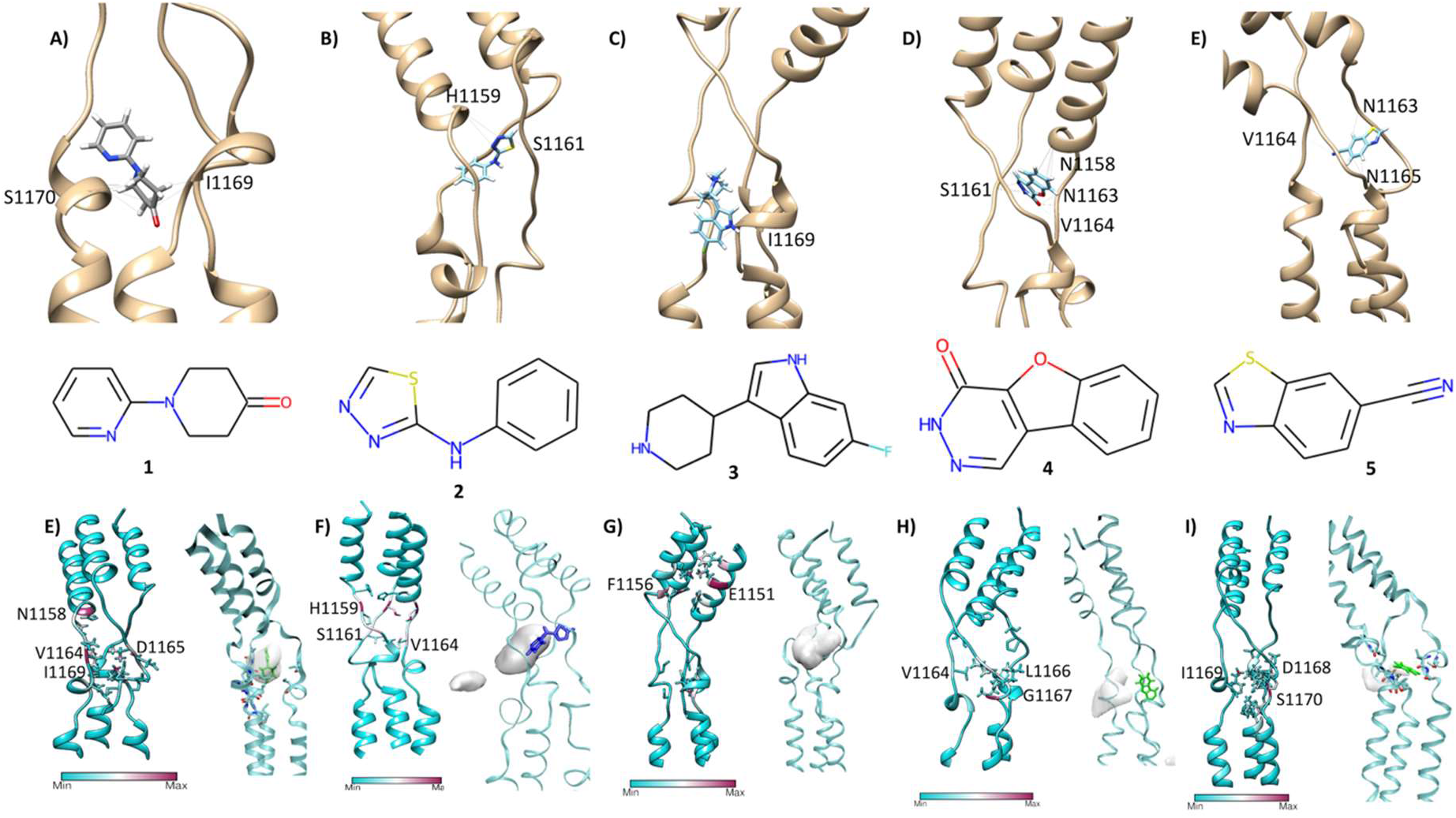
**A-D)** The best five fragments (1-5) according to molecular docking followed by 10 ns of post-docking cMD simulations; the final conformation (stick representation) after 10 ns is reported. **E-I**) Stalk residues (ribbon) that most interacted with fragments 1-5 during 500 ns of cMD; residues with the highest occupancy are in maroon. The density maps (iso surface 20%, grey surfaces) of the fragments are also reported.

**Table 1:** Summary of the best 5 fragments after 500 ns of cMD.

| Fragment | IUPAC NAME | Ligand Position on the stalk before unbinding | Displacement Time |
| --- | --- | --- | --- |
| 1 | 1-pyridin-2-ylpiperidin-4-one | N-terminal | ~160 ns |
| 2 | N-phenyl-1,3,4-thiadiazol-2-amine | N-terminal | ~200 ns |
| 3 | 6-fluoro-3-piperidin-4-yl-1H-indole | N-terminal | ~300 ns |
| 4 | 3H-[1]benzofuro[2,3-d]pyridazin-4-one | C-terminal | ~200 ns |
| 5 | 1,3-benzothiazole-6-carbonitrile | C-terminal | ~110 ns |

### Mix MD to check the fragments’ accessibility to Conserved Region 1

Since mixMD simulations of the molecular probes BENZ, FAC, and MTA (**Figure S2**) suggested some degree of accessibility to the stalk despite the high glycosylation of the SP, we run further mixMD simulations using **1-3** (**Table 1, Figure 4**) to investigate the accessibility of larger compounds, and any convergence with the metastable configurations sampled during the 500 ns post-docking cMD simulations. During mixMD simulations, the fragments were able to reach the stalk protein surface on isolated spots, overcoming the shield provided by the glycans (**Figure S4**). Fragments **1** made fewer interactions with the stalk among the three compounds (**Table S4**), mainly engaging residues F1148, T1155, K1149, E1151, and L1152 **(Figure S4A, Video S2)**. Fragments **2** formed interatomic contacts with residues K1149, Y1155, E1151, L1152, V1176, H1159, and F1148 (**Table S4, Figure S4B**). Compounds **3** formed the most persistent interactions with the stalk, engaging Y1155, E1182, E1151, N1178, Q1180, K1149, H1159, L1152, R1185, ASP1153, F1156, V1177, and D1153 side **(Table S4, Figure S4C)**. Many of these residues were engaged with a high turnover by different fragment molecules, indicating low stability of the interactions. However, mixMD simulations indicated two cryptic binding sites where single molecules of **1-3** resided for almost the whole duration of the simulation. In sub-pocket 1 (**Figure 5B**), located in the Conserved Region 1, fragment **3** formed hydrogen bonds with E1151 and hydrophobic contacts with residues F1148, K1149, L1152, and Y1155. In sub-pocket 2 (**Figure 5C**) at the interface between Conserved Regions 1 and 2, fragment **3** formed hydrophobic contacts with L11,86, V1189, A1190, and K1191, and three hydrogen bonds with E1182, N1194, and N1187; glycan residues participated in the stabilization of the molecule through van der Waals contacts. Notably, N1187 is a Valine in the Delta strain, therefore, it is not conserved.

**Figure 5.**
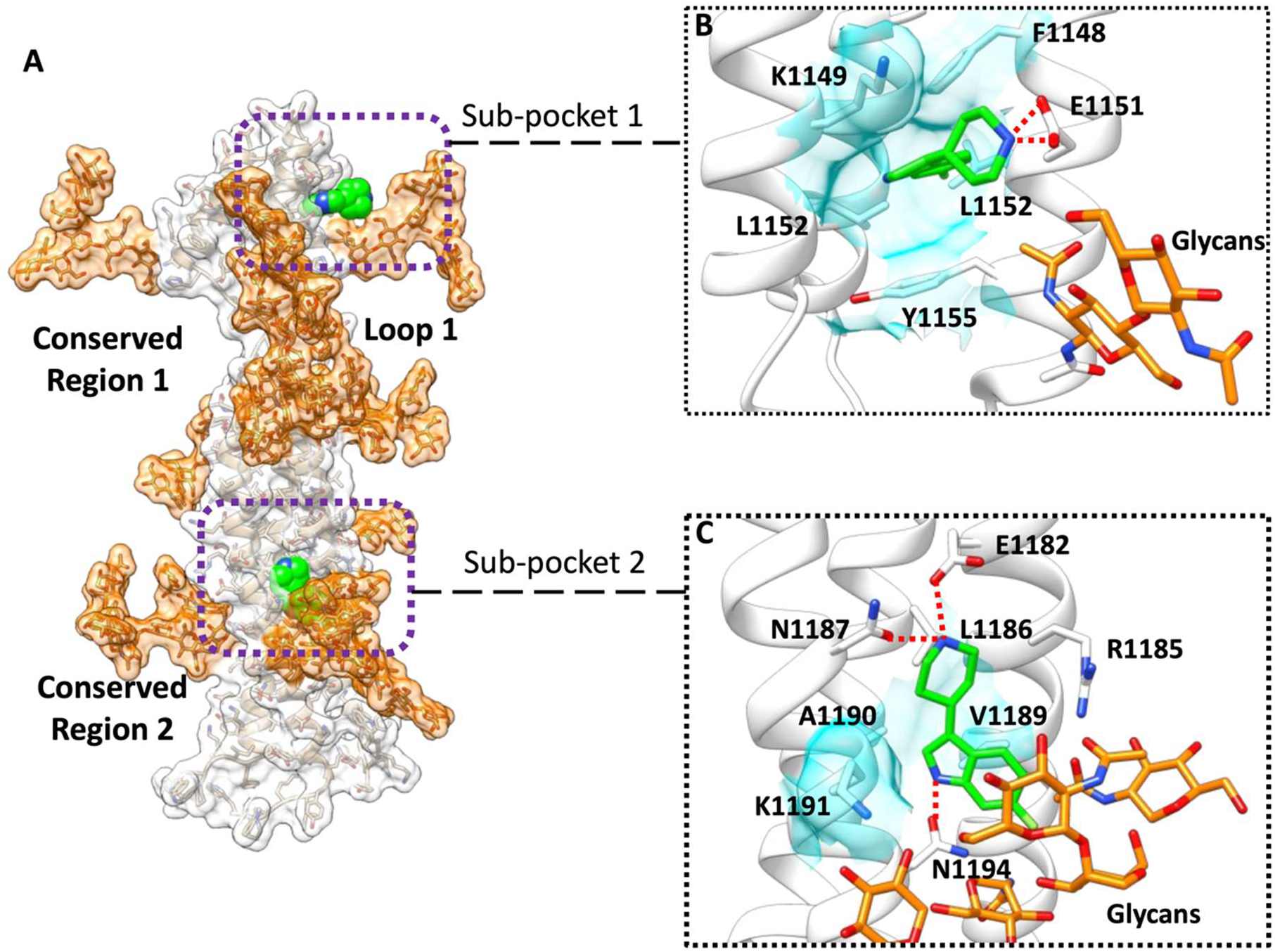
MixMD simulations indicated two sub-binding pockets. **A)** Overview of the stalk SP (white transparent surface); glycan residues are depicted in orange transparent surface, while fragment **3** is in green van der Waals spheres. **B**) magnification of the sub-pocket 1; **C**) magnification of the sub-pocket 2; fragment **3** is in green stick representation, while glycan residues are orange. Hydrogen bonds are shown ad dashed red lines.

## Conclusion

The SP stalk region is conserved amongst the SARS-CoV-2 VOCs. Given its role in orienting the RBD for binding to ACE2, impairing the flexibility of the loops formed by residues T1160-S1170 through the binding of a drug, could represent a therapeutic approach to explore^38,64^. We investigated the druggability of the SP stalk using 240 molecular fragments and considered the shielding effect of the glycans on the protein surface. Our computational workflow combined molecular docking, high-throughput MD simulations, and mixMD as orthogonal methods to evaluate putative interactions on the stalk region. Molecular docking predicted putative interaction sites around residues T1160 - S1170. High throughput cMD simulations of 559 docking poses suggested the instability of docking poses, except for a few ligands that were then further evaluated in longer simulations. Metastable interactions on the whole stalk were confirmed in the proximity of residues H1159 – I1169. Finally, mixMD simulations of the three most promising fragments **1-3** sampled two narrow binding sites within the helices of the stalk, which allowed for binding. Sub-pocket 1 is located just nearby Loop1 and might represent an anchor point for the design of larger ligands bearing a group that intercalates between the stalk helices and a flexible moiety that impairs Loop1 dynamics and likely the whole SP flexibility, although this strategy appears complicated. Sub-pocket 2 presents a balanced mix of hydrophilic and hydrophobic residues that increase the druggability compared to sub-pocket 1. However, it bears the not-conserved residue N1147, it is more distant from Loop 1 than pocket 1, and it is heavily influenced by glycan residues. In summary, our work highlights the challenges of exploiting the SP stalk as a therapeutic target. A fragment-based approach appears not suitable for this task. Future alternative studies could be based on the symmetry of the trimeric SP stalk and higher molecular weight ligands.

## Supporting information

Supplementary information, figures and graphs

Fragment 2

Fragment 1

## ACKNOWLEDGMENT

GD is grateful to SilcSBio for the provision of a local version of CGenFF, which was used to automatize the parameterization of the compounds.

