## Supplementary information, figures and graphs for "Is the Stalk of the SARS-CoV-2 Spike Protein Druggable?"

Centre for Sport, Exercise and Life Sciences (CSELS), Faculty of Health and Life Sciences, Coventry University, Coventry CV1 2DS, UK

**Video S1. Fragment 1-HR1 Binding and Displacement.** Fragment **1** docked pose is between the three HR1 chains. The motion of the chains leaves enough space for the fragment to rotate and move, indicating a weak affinity. The initial binding seems to be due to the steric hindrance by the three chains. After about 200ns, a conformational change of the loops opens a path for the fragment to unbind. Comparable results were obtained with fragments **2**, **3**, **4**, **5**. The density map of fragment **1** during the simulation (iso value 20%) is shown.

**Video S2. Fragment 2-HR1 mixMD simulation.** Fragment **1** identifies three interacting sites: one at the base of the HR1, one at the center of the flexible loop, and one on top of the HR1, on the terminal portion of the domain. Contacts were transitory and not indicative of a strong binding. However, mixMD identified a few stable niches along the stalk axis alongside the N-Terminal region and below the flexible loops. We focused on the conserved sequences between all SARS-CoV-2 variants and, therefore, discarded other results. Comparable results were obtained for fragments **2**, **3**, **4**, and **5**. The density map of **1** during the simulation (iso value 5%) is shown.

**Commented [GD1]:** please change frags codes to number 1 to 5 as per main text

**Table S1. SMILES of the fragments used**.

**Table S2. Fragments' docking poses ranked according to the docking score.**

**Table S3. DES of the 559 fragments.** The average (avg) and standard deviation (SD) of DES, GBSA, and RMSD are also reported.

**Table S4. The best 18 fragments according to post-docking MD**.

**Table S5. The best 5 fragments (**1-pyridin-2-ylpiperidin-4-one, N-phenyl-1,3,4-thiadiazol-2-amine, 6-fluoro-3-piperidin-4-yl-1H-indole); their contacts' occupancies are indicated individually with the respective frequency expressed in % over the totality of the MD frames.

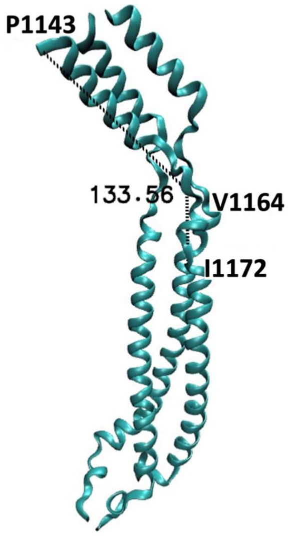


**Figure S1.** Our simulation indicated that an angle of 133.56° is formed between P1143, V1164, and I1172 causing important changes in the structure with the consequent opening of broad gaps between the chains with P1143 and G1171 being key residues for structural flexibility.

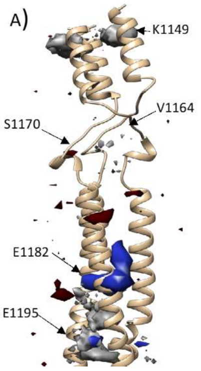


**Figure S2.** MixMD Density maps for BENZ (grey), FAC (blue), and MTA (red) at the 20% iso value. The probes identified possible sites on the stalk (ribbon) accessible to small molecules, despite the presence of glycans (not shown for clarity).

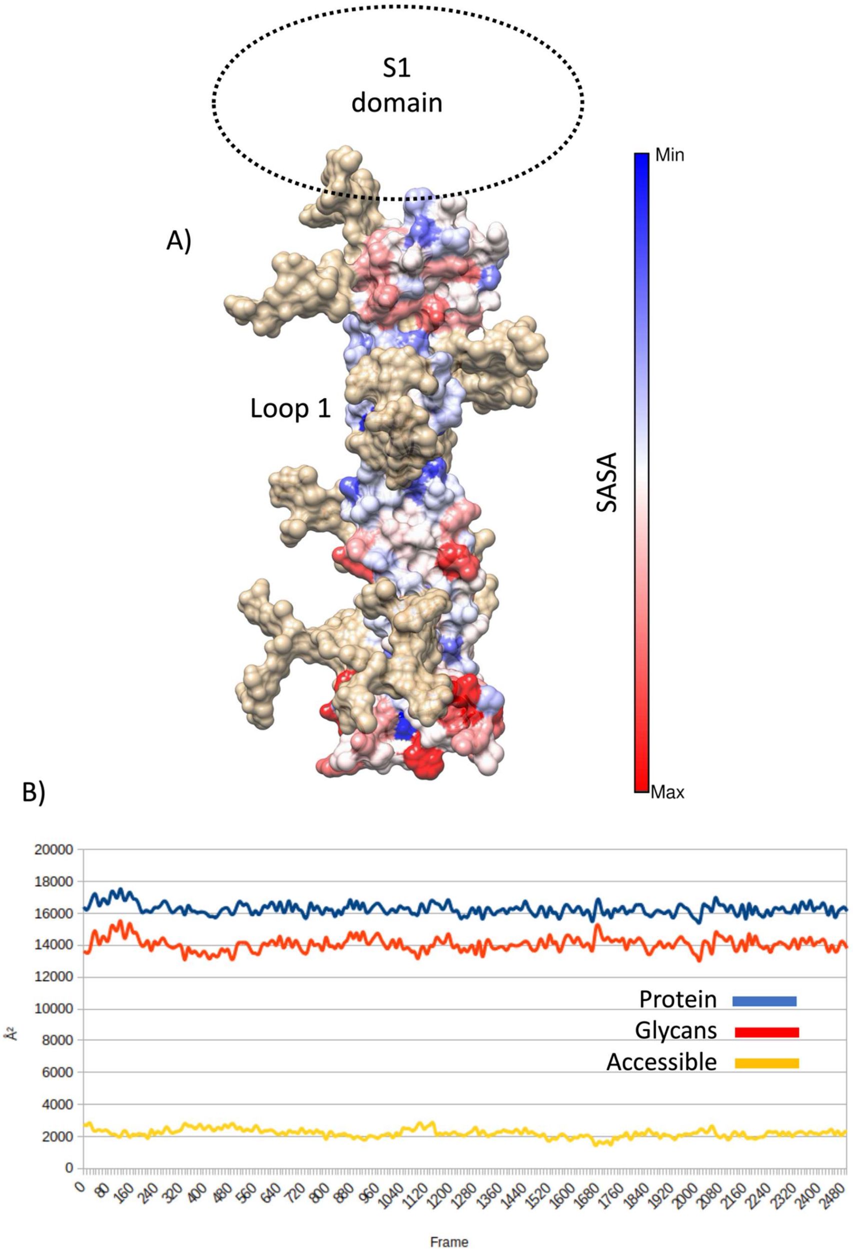


**Figure S3. A)** Solvent Accessible Surface Area (SASA) indicates possible accessible sites (red color) toward the N-terminal of the stalk, below the connection with S1, and below the flexible Loop 1; glycans are shown as tan surface. **B**) Glycans' movement tends to give access to a small portion of the stalk and has a considerable impact on the accessible surface.

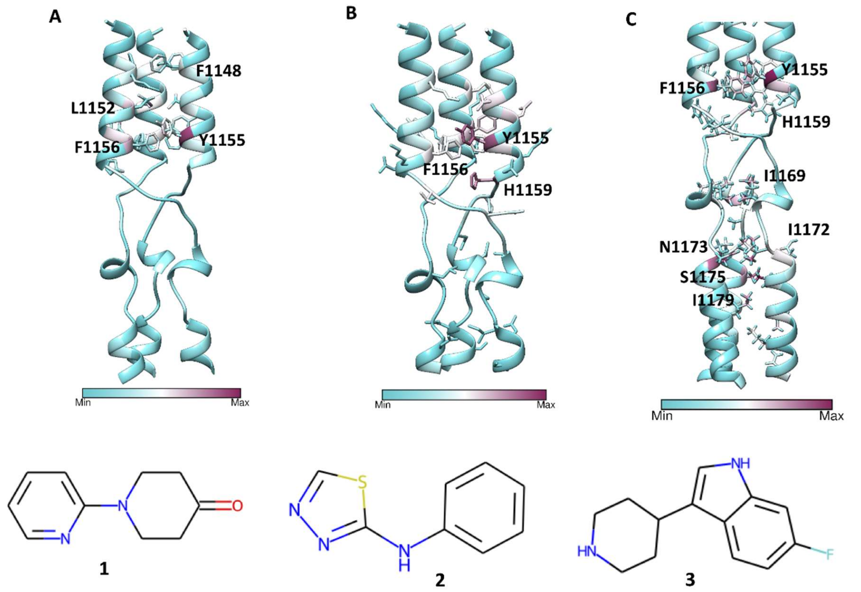


**Figure S4. Contacts formed during mixMD simulations of fragments 1-3.** The stalk is reported as a cyan ribbon, while the most involved residues (**Table S5**) are in stick maroon representation.
